# Assessing the Sylvatic Yellow Fever Vectors in Southern Brazil

**DOI:** 10.1101/2025.06.12.659296

**Authors:** Sabrina F. Cardoso, Larissa Akemi Oliveira Kikuti, Andre Akira Gonzaga Yoshikawa, Iara Carolini Pinheiro, João Victor Costa Guesser, Maycon Sebastião Alberto Santos Neves, Dinair Couto-Lima, Renata Rispoli Gatti, Josiane Somariva Prophiro, André N. Pitaluga, Luísa D. P. Rona

## Abstract

Yellow fever (YF) is an infectious disease caused by the yellow fever virus (YFV), an arbovirus of the *Flaviviridae* family. It is transmitted through the bite of infected mosquitoes belonging to the Culicidae family and affects both humans and non-human primates (NHPs).

This study aimed to investigate the sylvatic Culicidae fauna and the occurrence of natural YFV infection in a microregion of southern Santa Catarina, Brazil, an area recently affected by a sylvatic YF outbreak. Entomological collections were carried out between January and February 2023 in five municipalities with confirmed viral circulation.

Natural YFV infection was assessed using RT-LAMP. A total of 4,352 female culicids were collected, representing at least 32 species, including several key sylvatic YFV vectors. *Haemagogus leucocelaenus* was identified in all sampled municipalities, whereas *Haemagogus janthinomys*, the primary vector of sylvatic YFV in Brazil, was not detected.

Mosquitoes from the genera *Aedes*, *Haemagogus*, *Psorophora*, and *Sabethes* were tested for YFV. Only one pool, composed of *Sabethes albiprivus*, tested positive, yielding a minimum infection rate (MIR) of 11.6. This is the first record of natural YFV infection in *Sa. albiprivus* in southern Brazil, and only the third reported case globally, highlighting its potential role as a secondary vector in maintaining viral circulation in sylvatic environments.

Based on species presence and abundance, *Hg. leucocelaenus* is likely to have acted as the primary YFV vector in the study area. The composition of the culicid fauna, coupled with the detection of YFV in sylvatic vectors, indicates an ongoing epidemiological risk. These findings underscore the need to strengthen entomological surveillance and expand YF vaccination coverage in affected and neighbouring regions.

**Author Summary:** This study investigated sylvatic mosquito populations in a region of southern Santa Catarina, Brazil, recently affected by a yellow fever outbreak. Yellow fever is a serious mosquito-borne disease that can affect both humans and non-human primates. Nearly 4,400 mosquitoes from various species were collected, and, for the first time in southern Brazil, natural infection with yellow fever virus (YFV) was detected in *Sabethes albiprivus*. This finding suggests that *Sa. albiprivus* may play a previously unrecognized role in maintaining YFV in the environment.

The known vector *Haemagogus leucocelaenus* was found in all sampled locations, indicating it may have been the primary vector responsible for virus transmission in the region. These results enhance our understanding of YFV’s natural transmission cycles and provide new insights into how the virus persists in sylvatic environments.

This research contributes to the fields of biology, ecology, and public health by reinforcing the importance of ongoing entomological surveillance and preventive vaccination, both of which are essential for preventing future outbreaks and protecting vulnerable populations.

## Introduction

The Culicidae family comprises a large and widespread group of mosquitoes, including more than 3,700 species across 113 genera, found in both temperate and tropical regions worldwide [1]. Approximately 150 of these species are known vectors of humans diseases, significantly contributing to global morbidity and mortality. One such disease is yellow fever (YF), caused by the yellow fever virus (YFV) [2, 3], a member of the *Flaviviridae* family and classified under the species *Orthoflavivirus flavi* [4].

Yellow fever is endemic in sub-Saharan Africa, Central and South America, and the Caribbean [5], causing thousands of cases and deaths annually in Africa and South America, despite the availability of effective vaccines [6]. In Brazil YF is classified into two distinct transmission cycles - sylvatic and urban - which differ significantly in terms of vector species, vertebrate hosts, and geographic distribution [3]. There have been no reported cases of the urban transmission cycle in the country since 1942 [6]. However, in 2016, the virus re-emerged, triggering the most severe sylvatic YF outbreak in the past 80 years. This outbreak affected both humans and non-human primates (NHPs) [7, 8]. The virus initially spread from Venezuela into the Brazilian Amazon, then advanced through the Centre-West and Southeast regions, reaching the Atlantic Forest coastal areas and rapidly extending southward, where it remained active at least until the end of 2021 [8, 9].

*Aedes* (*Stegomyia*) *aegypti* is the primary vector responsible for YFV transmission in the urban cycle, while species from the genera *Haemagogus* and *Sabethes* transmit the virus in the sylvatic cycle as primary and secondary vectors, respectively [6]. In the Americas, the most important sylvatic vector is *Haemagogus janthinomys*, which is widely distributed throughout Brazil [10]. Other *Aedes* species, beyond *Ae*. *aegypti*, have also been found naturally infected with the YFV [6, 11]. Furthermore, mosquitoes of the genus *Psorophora* have been identified as natural hosts of the virus [3, 12], indicating their potential role in transmission.

Several studies have investigated the potential YFV vectors in Brazil [13, 6, 14, 15, 16]. However, there are few records of the sylvatic mosquito fauna in the state of Santa Catarina, southern Brazil. Therefore, the present study aimed to identify sylvatic mosquito species in a microregion of southern Santa Catarina that was recently affected by a sylvatic YF outbreak. Additionally, we investigated natural YFV infection in potential vector species to better understand their role in the transmission and spread of the outbreak.

## Materials and methods

### Ethics statement

All members of the research team were vaccinated against YFV prior to the commencement of the study. To minimise the risk of exposure to pathogens during fieldwork, the team also wore personal protective equipment, including long-sleeved clothing, caps, and boots.

### Study area

Entomological surveys were conducted in a microregion located in the southern part of Santa Catarina state, Brazil (Fig 1). The area has a temperate tropical climate and contains remnants of Atlantic Forest vegetation [17]. In February 2021, YFV circulation was confirmed in the region for the first time through the detection of infected NHPs [18]. As a result, mosquito collections were carried out in municipalities with confirmed viral circulation: Santa Rosa de Lima (−28.032639, −49.149684), Rio Fortuna (−28.141177, - 49.146304), Braço do Norte (−28.195246, −49.136690), Pedras Grandes (−28.514485, - 49.242145), and São Martinho (−28.127728, −49.051721) (Fig 1).

**Fig. 1.**
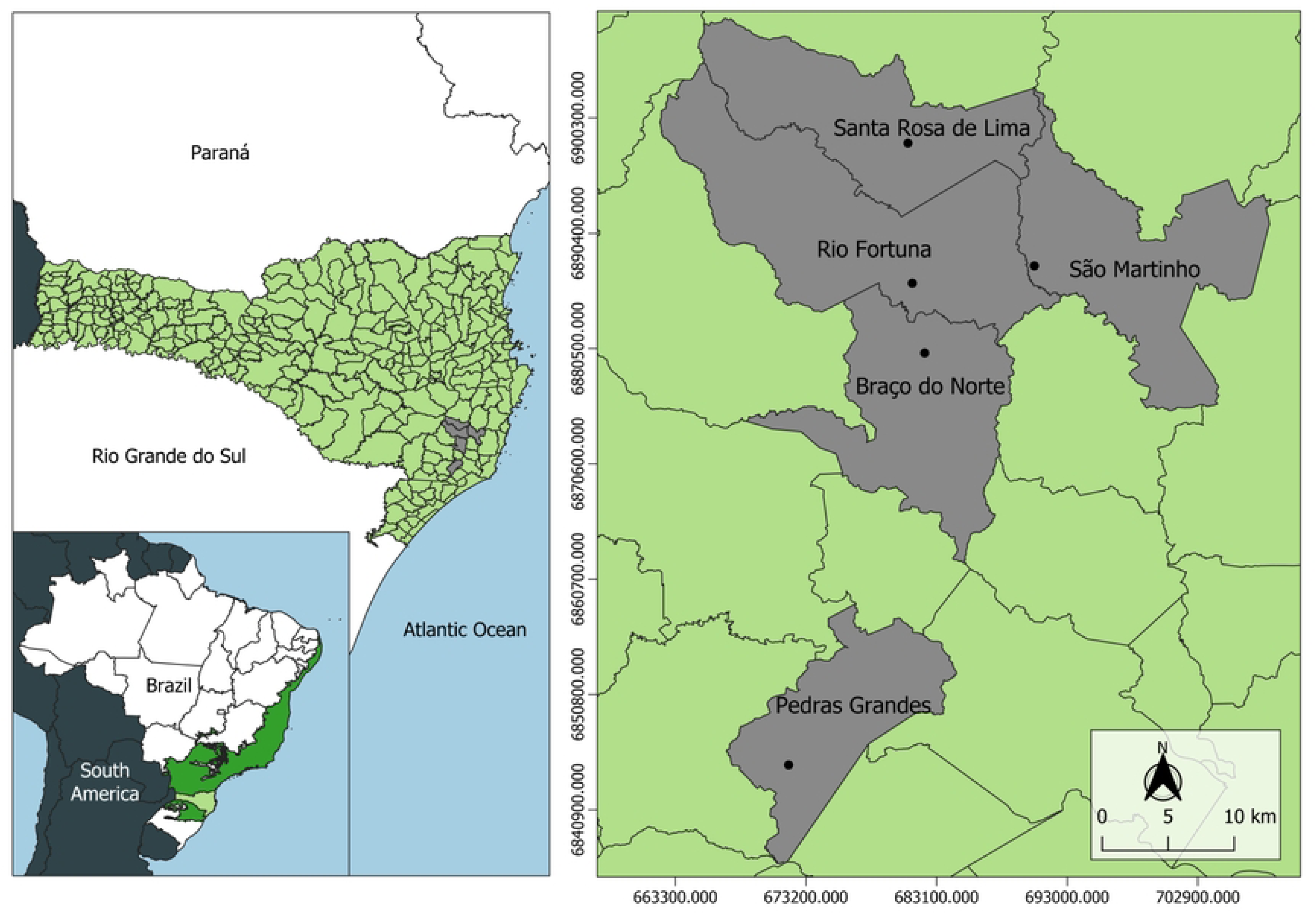
Geographic distribution of sylvatic mosquito collection sites in municipalities of southern Santa Catarina State, Brazil. The map highlights the state of Santa Catarina in light green, with the original Atlantic Forest cover in Brazil indicated in dark green. Municipalities included in the study are marked in grey. The map was generated using Quantum GIS (QGIS, version 3.34) [19].

### Entomological collection and species identification

Adult female mosquitoes were collected during the summer of 2023, specifically in January and February. Sampling was conducted between 08:00 and 17:00 on three consecutive days in each of the five municipalities. In each of the 15 collections, mosquitoes were captured simultaneously in both the tree canopy and at ground level using an entomological net and a Castro-type oral aspirator. Additionally, four CDC light traps (CDC-LT) baited with CO_2_ (dry ice) were installed in the canopy. Following collection, specimens were stored and transported on dry ice.

In the laboratory, mosquitoes were morphologically identified using a stereomicroscope (SZX16 Olympus) on a cold surface. Identification to genus and species level was carried out using standard taxonomic keys for culicids [10, 20, 21, 22]. Specimens belonging to the genera *Haemagogus*, *Sabethes*, *Aedes*, and *Psorophora* were grouped into pools of up to 10 individuals, based on species and collection site, and stored at −80°C for subsequent viral detection.

### Standardisation of viral RNA extraction

RNA extraction was standardised based on a modified protocol described by Silva et al. [23]. To simulate viral infection, pools of up to 10 mosquitoes were homogenised in 250 µL of nuclease-free water (UltraPure™, Invitrogen, cat. no. 10977015) containing 2.5 µL of RNAsecure (Invitrogen, cat. no. AM7006) and spiked with 2.5 µL of total RNA (80 ng/µL) extracted from Vero cells infected with the YFV 17D vaccine strain. After brief centrifugation, 140 µL of the supernatant was used for RNA extraction with the QIAamp Viral RNA Mini Kit (Qiagen, cat. no. 52904), following the manufacturer’s instructions. To validate the RNA extraction, the samples were tested with RT-LAMP assays (data not shown). Further details about the RT-LAMP assays can be found in the *Molecular identification of YFV in mosquitoes* section.

The same RNA extraction procedure was applied to mosquito pools, each with up to 10 individuals from the genera *Haemagogus*, *Sabethes*, *Aedes*, and *Psorophora*, to detect natural YVF infection.

### Molecular identification of YFV in mosquitoes

To detect YFV in captured sylvatic mosquitoes, RNA extracted from mosquito pools was used in a colorimetric RT-LAMP assay, performed with the 2X WarmStart Colorimetric LAMP Master Mix kit (New England BioLabs, Protocol M1800). The assay targeted the NS1, NS5, and E genes of the viral genome, using a modified protocol based on Cardoso et al. [18], with an optimised incubation time of 60 minutes. Results were interpreted visually: a pink colour indicated a negative result, while yellow indicated a positive result. Reactions were documented with a smartphone camera.

### Data analysis

Quantitative and qualitative analyses of the mosquito fauna were conducted using Microsoft® Excel® (Microsoft 365 MSO: version 2504; Build 16.0.18730.20122). To assess species dominance and distribution, the relative abundance of each species was calculated as the proportion of individuals of a given species relative to the total number of mosquitoes collected.

Assuming each positive pool contained only one infected mosquito, the minimum infection rate (MIR) for YFV was determined by dividing the number of positive pools by the total number of mosquitoes of that species tested and multiplying the result by 1000 [6].

## Results and discussion

### Abundance and diversity of sylvatic culicid fauna

This is the first study to survey sylvatic mosquito fauna along the southern coast of Santa Catarina, Brazil. Over the study period, 15 mosquito collections were conducted across five municipalities, yielding 4352 female specimens. At least 32 taxa were identified to species level and 14 to genus level, indicating high species diversity (Table 1). The sample included genera of recognised epidemiological importance, such as *Aedes*, *Anopheles*, *Haemagogus*, and *Sabethes*, highlighting public health relevance, as vector composition and diversity strongly influence the risk of pathogen transmission [24]. The most frequent species were *Trichoprosopon townsendi* (21%), *Limatus durhamii* (9%), *Aedes scapularis* (5), and *Psorophora ferox* (4%).

**Table 1.**
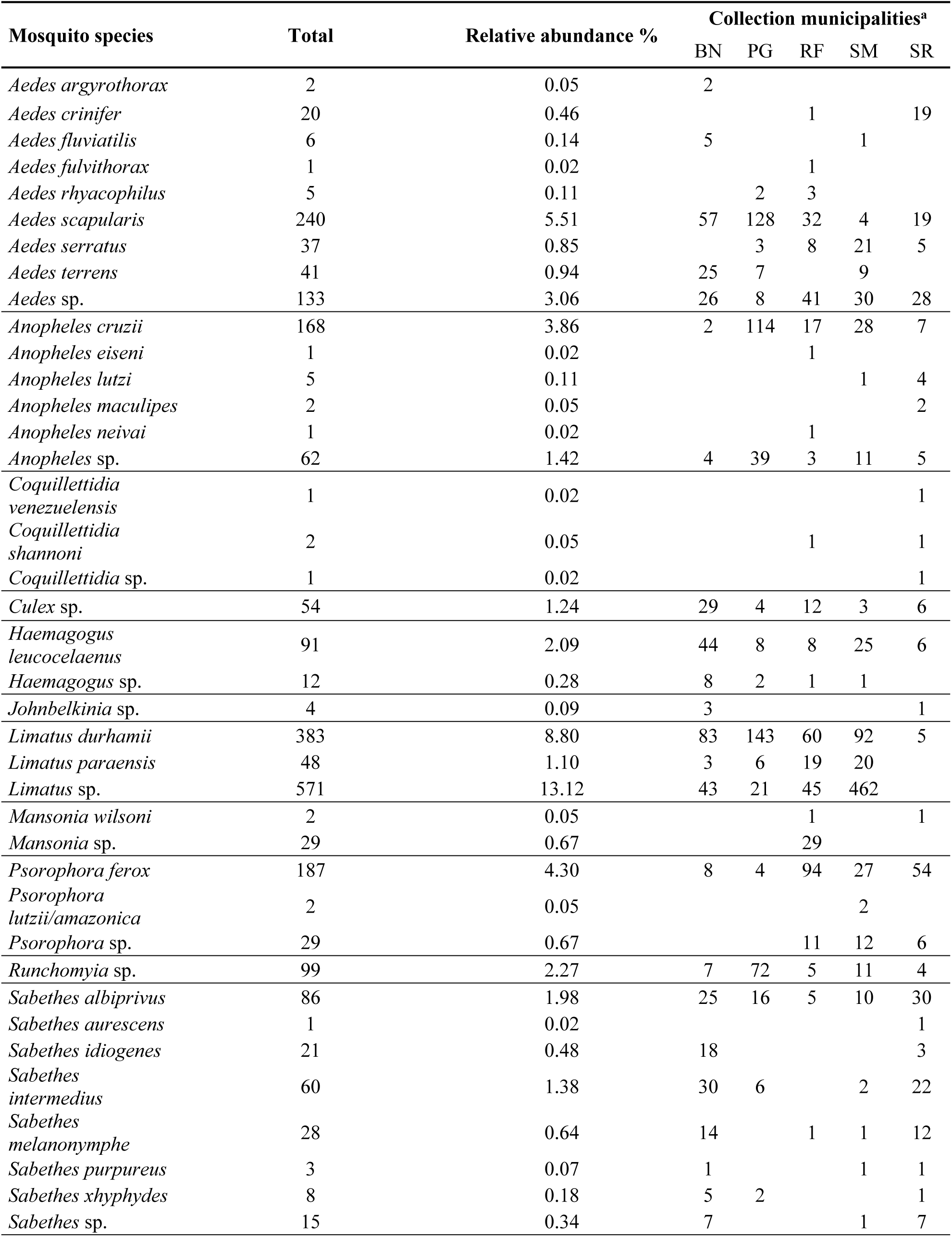
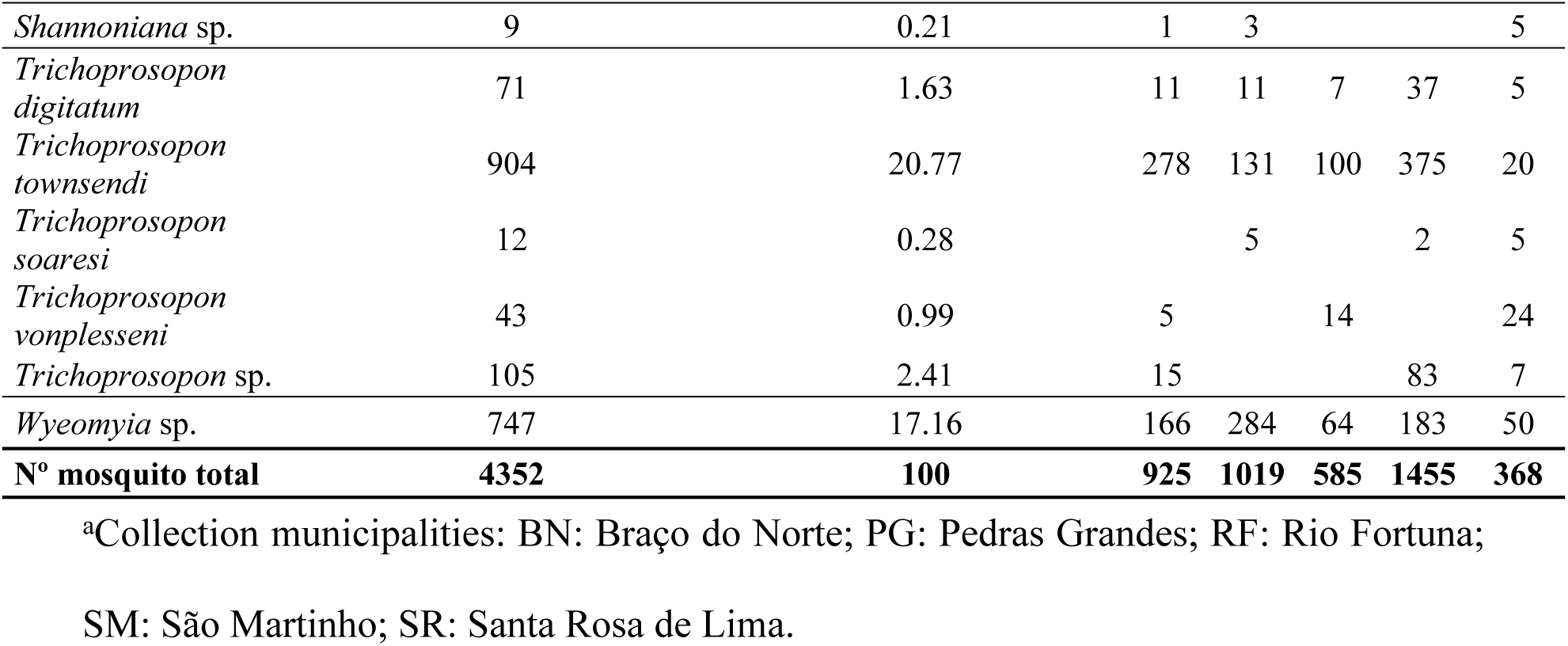
Adult mosquito species, grouped by genus, collected from January to February 2023 in five municipalities of Santa Catarina State, southern Brazil.

The mosquito abundance recorded in this study is consistent with the findings of Deus et al. [25], who also reported *Wyeomyia* and *Limatus* among the most frequently collected genera in São Paulo State. In our data, *Wyeomyia* comprised 17% of specimens and was the most abundant genus in Pedras Grandes (284 individuals), while *Limatus* accounted for 13% and was dominant in São Martinho (462 individuals) (Table 1). These were the second and third most widespread genera observed overall.

A notable exception was *Tr. townsendi*, the most abundant species in our study, comprising 21% of all specimens. It was particularly common in Braço do Norte and Rio Fortuna, where 278 and 100 individuals were recorded, respectively. However, this species was not found in the survey by Deus et al. [25] in São Paulo State.

In contrast, our findings align with those of Orlandin et al. [26], who reported *Trichoprosopon pallidiventer* as the most common species in western Santa Catarina, accounting for over 59% of nearly 1000 specimens. These results suggest that *Trichoprosopon* species can reach high local abundances due to their strong ecological adaptability and resilience, which support their successful establishment across diverse habitats [27].

*Aedes scapularis*, *Li. durhamii*, and *Ps. ferox* are frequently reported in high numbers in entomological surveys [6, 13]. For instance, a study conducted in southeastern Brazil, including the states of Rio de Janeiro, São Paulo, Espírito Santo, and Minas Gerais, identified *Ae. scapularis* as the most abundant species, representing 19% of more than 15000 mosquitoes collected [6]. In the same study, *Li*. *durhamii* and *Ps. ferox* ranked as the sixth and the eighth most collected species among over 80 recorded taxa, further highlighting their widespread occurrence. This pattern aligns with our findings, in which *Li*. *durhamii*, *Ae. scapularis* and *Ps. ferox* accounted for 9% (383 specimens), 5% (240 specimens), and 4% (187 specimens) of the total catch, respectively, making them the second, third, and fourth most abundant species in our survey.

Mosquitoes of the genus *Haemagogus* accounted for 2.4% of all specimens collected (103 individuals), with *Haemagogus leucocelaenus*, a primary sylvatic YF vector, representing 2.1% (91 individuals). This species was captured at both canopy and ground levels across all five municipalities (S1 Table), but was more frequently found in the canopy, consistent with its primatophilic behaviour and known preference for feeding in the forest canopy [6, 28].

The genus *Sabethes* comprised 5.1% of the total collection (222 individuals). The most frequently encountered species within this group was *Sabethes albiprivus*, a recognized secondary vector of sylvatic YFV, accounting for 2% of specimens (86 individuals), and recorded in all sampled municipalities. This aligns with the findings of Abreu et al. [6], who reported *Sa. albiprivus* as comprising 3% of the mosquitoes collected in a similar study conducted in southeastern Brazil.

### Natural infection of mosquitoes with yellow fever virus

In Brazil, most YFV isolates from mosquitoes have been obtained from species in the *Haemagogus* and *Sabethes* genera. However, natural infections have occasionally been reported in *Aedes* [3, 13] and *Psorophora* species [3, 12]. Based on this, we tested specimens from these four genera for YFV using RT-LAMP, analysing 154 pools in total (Table 2): 61 *Aedes*, 29 *Haemagogus*, 28 *Psorophora*, and 36 *Sabethes*.

**Table 2.** Numbers of adult mosquitoes from the *Sabethes*, *Haemagogus*, *Aedes*, and *Psorophora* genera tested for YFV infection, grouped by the number of pools tested, showing the number of positive pools and the corresponding minimum infection rates (MIR).

| Mosquito species | N° mosquitoes | Pools tested (positives) | MIR |
| --- | --- | --- | --- |
| *Sabethes albiprivus* | 86 | 10 (1) | 11.63 |
| *Sabethes intermedius* | 60 | 8 (0) | 0 |
| *Sabethes melanonymphe* | 27 | 5 (0) | 0 |
| *Sabethes idiogenes* | 21 | 3 (0) | 0 |
| *Sabethes xhyphydes* | 8 | 3 (0) | 0 |
| *Sabethes purpureus* | 3 | 3 (0) | 0 |
| *Sabethes aurescens* | 1 | 1 (0) | 0 |
| *Sabethes* sp. | 15 | 3 (0) | 0 |
| *Haemagogus leucocelaenus* | 91 | 24 (0) | 0 |
| *Haemagogus* sp. | 12 | 5 (0) | 0 |
| *Aedes argyrorhox* | 2 | 1 (0) | 0 |
| *Aedes crinifer* | 20 | 3 (0) | 0 |
| *Aedes fluviatilis* | 6 | 2 (0) | 0 |
| *Aedes fulvithorax* | 1 | 1 (0) | 0 |
| *Aedes rhyacophilus* | 5 | 2 (0) | 0 |
| *Aedes scapularis* | 240 | 26 (0) | 0 |
| *Aedes serratus* | 37 | 6 (0) | 0 |
| *Aedes terrens* | 41 | 5 (0) | 0 |
| *Aedes* sp. | 133 | 15 (0) | 0 |
| *Psorophora ferox* | 187 | 21 (0) | 0 |
| *Psorophora lutzii/amazonica* | 2 | 1 (0) | 0 |
| *Psorophora* sp. | 29 | 6 (0) | 0 |
| **Total** | **1027** | **154** | **11.63** |

Among the 36 *Sabethes* pools, only one tested positive for YFV, pool 32, composed of *Sa. albiprivus* specimens (Figs 2 and 3). This result was confirmed by three independent retests (S1 Fig). The positive pool contained 10 mosquitoes collected in the municipality of São Martinho, corresponding to a MIR of 11.63. No YFV-infected mosquitoes were detected in the remaining pools, including the 24 pools *of Hg. leucocelaenus* (Table 2), a species traditionally recognised as a YF vector [13].

**Fig 2.**
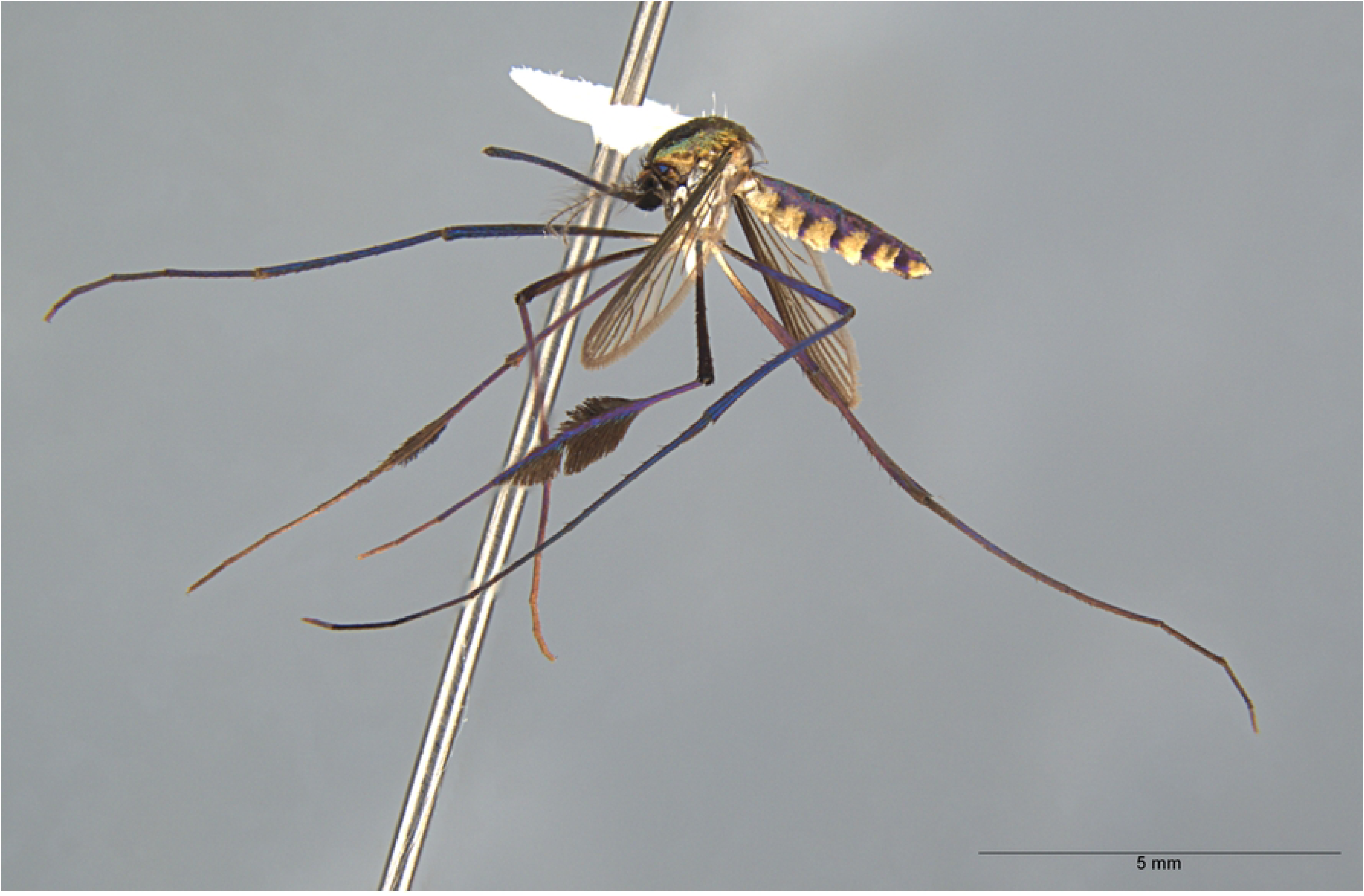
Image of *Sabethes albiprivus*, a specimen from the Fiocruz-CCULI collection. Fiocruz-CCULI archive.

**Fig 3.**
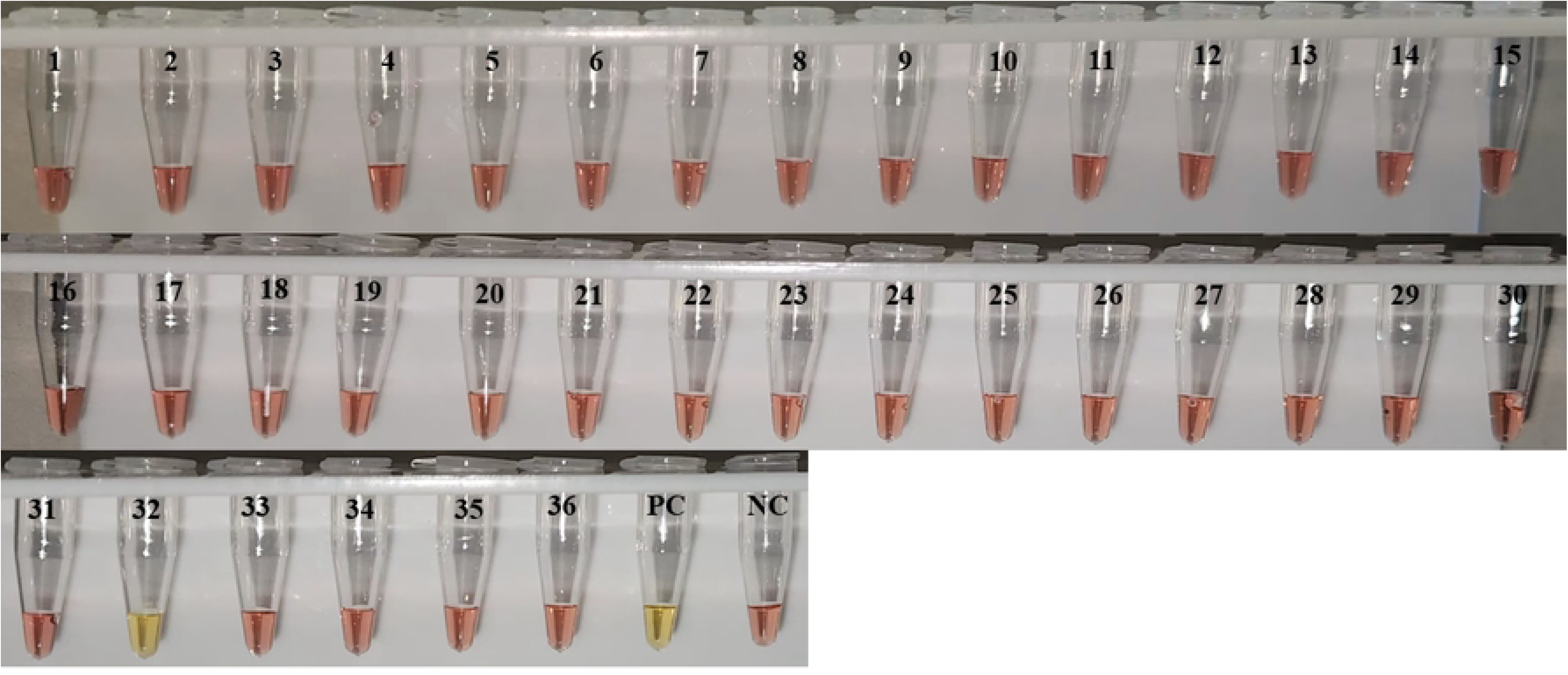
RT-LAMP analysis of mosquito pools from the genus *Sabethes*. RT-LAMP reactions were performed at 65 °C for 60 minutes. Yellow indicates positive results; pink indicates negative. Tubes 1–31 and 33–36 correspond to *Sabethes* pools negative for YFV. Tube 32, composed of *Sa. albiprivus* specimens, tested positive for YFV. NC = negative control (nuclease-free water); PC = positive control (YFV RNA, vaccine strain 17D).

*Sabethes albiprivus* is widely distributed across Brazilian biomes [6, 15, 29], with confirmed records in the southern states of Paraná [30], Santa Catarina [31], and Rio Grande do Sul [11, 16]. YFV infected individuals have also been reported in Argentina [32] and in Minas Gerais, Brazil [15]. Our study presents the first confirmed natural YFV infection in *Sa. albiprivus* in southern Brazil, only the second such case in the country and the third globally, highlighting its potential epidemiological importance. In the Brazilian Cerrado, particularly in northern Minas Gerais, *Sa. albiprivus* has been recognized as a secondary vector, with a MIR of 3.3 [15]. In contrast, our study recorded a MIR of 11.6, nearly three times higher, suggesting a greater role in YFV circulation in southern Brazil. Its involvement in YFV maintenance and secondary transmission has previously been proposed [6, 15], and laboratory studies have confirmed its vector competence [33].

Rezende et al. [34] reported the introduction and persistence of YFV in southeastern Brazil from 2016 to 2018, emphasising that ecological and climatic conditions outside the Amazon Basin, such as those in the Atlantic Forest, can support viral maintenance during interepidemic periods. Our findings align with this, as YFV was detected in 2023, nearly two years after the 2021 sylvatic outbreak, indicating continued circulation despite the absence of active transmission. This persistence likely reflects complex vector-host dynamics and possible hidden transmission cycles that may remain undetected. Overall, the evidence suggests that *Sa. albiprivus* played a secondary role in YFV transmission and contributed to viral persistence in the region.

During the last sylvatic YF outbreak in Brazil, *Hg. leucocelaenus* was considered the primary YFV vector in Brazil due to its widespread distribution, abundance, and high natural infection rates [6]. This species has been found infected with YFV in several regions [6, 28], including the southern state of Rio Grande do Sul [16]. In our study, 103 *Haemagogus* specimens were collected, including 91 *Hg*. *leucocelaenus*, yet none tested positive for viral RNA, despite their presence across all five sampled municipalities in Santa Catarina affected by YFV outbreaks. Although *Hg. janthinomys* is traditionally recognized as the main sylvatic YFV vector in Brazil [6, 13, 20], its absence in the sampled areas suggests that *Hg. leucocelaenus*, alongside *Sa. albiprivus*, may have played a key role in sustaining transmission during the recent outbreak in Santa Catarina.

The five municipalities surveyed are small, with urban areas that directly bordering natural habitats. These regions are characterised by fragmented vegetation linked by ecological corridors, facilitating mosquito movement in search of blood meals [8, 20]. The high dispersal capacity of YFV vectors, even in partially deforested landscapes [20], combined with declines in non-human primate (NHP) populations due to habitat loss and YF-related mortality, reduces available blood sources and promotes mosquito dispersal [8, 14]. Under these conditions, the studied municipalities represent high-risk areas for YFV transmission. Accordingly, widespread yellow fever vaccination campaigns are strongly recommended in these and surrounding regions, given the continued circulation of YFV and its sylvatic vectors.

### Conclusion

This study represents the first survey of sylvatic mosquito fauna and natural YFV infection in the state of Santa Catarina. The detection of naturally infected *Sa. albiprivus* highlights its role as an important vector in the maintenance and transmission of the virus in transitional zones between sylvatic and human-modified environments.

The absence of *Hg. janthinomys* and the presence of *Hg*. *leucocelaenus* suggest a potential change in local vector dynamics, particularly in areas where NHP populations were severely impacted. This further reinforcing the role of *Hg*. *leucocelaenus* as a primary YFV vector.

Given the continued circulation of the virus and the presence of competent vectors, this study underscores the importance of sustained entomological surveillance and expanded vaccination coverage to prevent future outbreaks in the region.

## Acknowledgments

We would like to thank the Municipal Health Departments of Santa Rosa de Lima, Rio Fortuna, Braço do Norte, São Martinho, and Pedras Grandes for their support in the entomological collections. We also extend our gratitude to the Department of Microbiology, Immunology, and Parasitology at the Federal University of Santa Catarina (UFSC) for donating YFV RNA (vaccine strain 17D), which was used as a positive control in the experiments.

## Authors contribution

SFC drafted the manuscript and collected the mosquito samples. SFC, ANP, AAGY, ICP, JVCG, and LDRP contributed to data generation and analysis. Sample processing and molecular analyses were performed by SFC, AAGY, ICP, LAOK, and JVCG. Mosquito identification was carried out by LAOK, SFC, and MSASN. DCL, MSASN, RRG, and JSP provided essential support with project logistics. ANP and LDRP contributed to manuscript preparation through critical review and also served as the principal investigators, overseeing the study’s design and coordination. All authors read and approved the final version of the manuscript.

## Conflict of interests

The authors declare that they have no competing interests.

This paper is part of the Ph.D. thesis of Sabrina Fernandes Cardoso from the Graduation Program of Cell and Developmental Biology (PPGBCD) at the Biological Sciences Center (CCB) at the Federal University of Santa Catarina (UFSC), Florianópolis, Brazil.

## Supporting information

**S1 Table.** Mosquito species captured by municipality. Collection sites: Braço do Norte (BN), Pedras Grandes (PG), Rio Fortuna (RF), São Martinho (SM), Santa Rosa de Lima (SRL). Canopy-level collection: Specimens were captured using CDC-type traps baited with CO₂ (CDC), as well as manually using an entomological net and a Castro-type oral aspirator (NET). Ground-level collection: Mosquitoes were manually collected at ground level using an entomological net and a Castro-type oral aspirator.

**S1 Fig.** RT-LAMP assay performed on a *Sabethes albiprivus* positive pool for YFV. The reaction was performed in triplicate. Reaction temperature: 65 °C; reaction time: 60 minutes. Yellow indicates a positive result; pink indicates a negative result. R1–R3: replicates 1 to 3. NC: negative control (nuclease-free water); PC: positive control (YFV RNA, vaccine strain 17D).

## References

1. Harbach RE. Culicidae. Mosquito Taxonomic Inventory. 2025. Available from: https://mosquito-taxonomic-inventory.myspecies.info/simpletaxonomy/term/6045. (accessed on 05 April 2025). Accessed April 5, 2025.

2. Guedes, MLP. Culicidae (Diptera) no Brasil: Relações entre enfermidades, distribuição e enfermidades. Oecol. Aust. 2012;16(2):283–296.

3. Vasconcelos PF. [Yellow Fever]. Rev Soc Bras Med Trop. 2003;36(2):275–93.

4. Postler TS, Beer M, Blitvich BJ, Bukh J, de Lamballerie X, Drexler JF, et al. Renaming of the genus *Flavivirus* to *Orthoflavivirus* and extension of binomial species names within the family Flaviviridae. Arch Virol. 2023 Aug 10;168(9):224.

5. McArthur DB. Emerging Infectious Diseases. Nurs Clin North Am. 2019 Jun;54(2):297–311.

6. Abreu FVS, Ribeiro IP, Ferreira-de-Brito A, Santos AACD, Miranda RM, Bonelly IS, et al. Haemagogus leucocelaenus and Haemagogus janthinomys are the primary vectors in the major yellow fever outbreak in Brazil, 2016-2018. Emerg Microbes Infect. 2019;8(1):218-31.

7. Barrett ADT. The reemergence of yellow fever. Science. 2018 Aug 31;361(6405):847-

8. Possas C, Lourenço-de-Oliveira R, Tauil PL, Pinheiro FP, Pissinatti A, Cunha RVD, et al. Yellow fever outbreak in Brazil: the puzzle of rapid viral spread and challenges for immunisation. Mem Inst Oswaldo Cruz. 2018 Sep 3;113(10):e180278.

9. Giovanetti M, Pinotti F, Zanluca C, Fonseca V, Nakase T, Koishi AC, et al. Genomic epidemiology unveils the dynamics and spatial corridor behind the Yellow Fever virus outbreak in Southern Brazil. Sci Adv. 2023 Sep;9(35):eadg9204.

10. Forattini OP. Culicidologia médica: identificação, biologia e epidemiologia. vol. 2. São Paulo: EDUSP; 2002. 864 p.

11. Cardoso Jda C, de Almeida MA, dos Santos E, da Fonseca DF, Sallum MA, Noll CA, et al. Yellow fever virus in Haemagogus leucocelaenus and Aedes serratus mosquitoes, southern Brazil, 2008. Emerg Infect Dis. 2010 Dec;16(12):1918-24.

12. Moreno ES, Rocco IM, Bergo ES, Brasil RA, Siciliano MM, Suzuki A, et al. Reemergence of yellow fever: detection of transmission in the State of São Paulo, Brazil, 2008. Rev Soc Bras Med Trop. 2011;44(3):290-6.

13. Abreu FVS, Ferreira-de-Brito A, Azevedo AS, Linhares JHR, de Oliveira Santos V, Hime Miranda E, et al. Survey on Non-Human Primates and Mosquitoes Does not Provide Evidences of Spillover/Spillback between the Urban and Sylvatic Cycles of Yellow Fever and Zika Viruses Following Severe Outbreaks in Southeast Brazil. Viruses. 2020 Mar 26;12(4):364.

14. Pinheiro GG, Rocha MN, de Oliveira MA, Moreira LA, Andrade Filho JD. Detection of Yellow Fever Virus in Sylvatic Mosquitoes during Disease Outbreaks of 2017⁻2018 in Minas Gerais State, Brazil. Insects. 2019 May 10;10(5):136.

15. de Oliveira CH, Andrade MS, Campos FS, da C Cardoso J, Gonçalves-Dos-Santos ME, Oliveira RS, et al. Yellow Fever Virus Maintained by Sabethes Mosquitoes during the Dry Season in Cerrado, a Semiarid Region of Brazil, in 2021. Viruses. 2023 Mar 15;15(3):757.

16. Vasconcelos PF, Sperb AF, Monteiro HA, Torres MA, Sousa MR, Vasconcelos HB, et al. Isolations of yellow fever virus from *Haemagogus leucocelaenus* in Rio Grande do Sul State, Brazil. Trans R Soc Trop Med Hyg. 2003;97(1):60–2.

17. Da Silva DA, Pfeifer M, Pattison Z, Vibrans AC. Drivers of leaf area index variation in Brazilian Subtropical Atlantic Forests. Forest Ecology and Management. 2020; 476;118477.

18. Cardoso SF, Yoshikawa AAG, Pinheiro IC, Granella LW, Couto-Lima D, Neves MSAS, et al. Development and validation of RT-LAMP for detecting yellow fever virus in non-human primates samples from Brazil. Sci Rep. 2024 Sep 28;14(1):22520.

19. QGIS Development Team, 2023. QGIS Geographic Information System. Open Source Geospatial Foundation. [Online] Available at: https://qgis.org. Accessed: April 30, 2025.

20. Consoli RAGB, Lourenço-de-Oliveira R. Principais mosquitos de importância sanitária no Brasil. Rio de Janeiro: Fiocruz; 1994. 228 p.

21. Lane, J. Neotropical Culicidade; University of São Paulo Publisher: São Paulo, Brazil, 1953; Vol. 1.

22. Neves MSAS, Motta MA, Maciel-DE-Freitas R, Xavier ADS, Loureno-DE-Oliveira R, Silva-DO-Nascimento TF. Illustrated identification key to females of the genus Sabethes Robineau-Desvoidy recorded from Brazil (Diptera: Culicidae), in dichotomous and interactive formats, including an updated list of species and new records for the states. Zootaxa. 2024 Feb 5;5406(2):253–87.

23. Silva SJRD, Paiva MHS, Guedes DRD, Krokovsky L, Melo FL, Silva MALD, et al. Development and Validation of Reverse Transcription Loop-Mediated Isothermal Amplification (RT-LAMP) for Rapid Detection of ZIKV in Mosquito Samples from Brazil. Sci Rep. 2019 Mar 14;9(1):4494.

24. Keesing F, Holt RD, Ostfeld RS. Effects of species diversity on disease risk. Ecol Lett. 2006 Apr;9(4):485–98.

25. Deus JT, Mucci LF, Lucheta Reginatto S, Pereira M, Bergo ES, de Camargo-Neves VLF. Evaluation of Methods to Collect Diurnal Culicidae (Diptera) at Canopy and Ground Strata, in the Atlantic Forest Biome. Insects. 2022 Feb 16;13(2):202.

26. Orlandin E, Santos EB, Piovesan M, Favretto MA, Schneeberger AH, Souza VO, et al. Mosquitoes (Diptera: Culicidae) from crepuscular period in an Atlantic Forest area in Southern Brazil. Braz J Biol. 2017 Mar;77(1):60–7.

27. Wilkerson RC, Linton Y-M, Strickman D. Mosquitoes of the World. Johns Hopkins University Press, Baltimore; 2021, Vol. 1 and Vol. 2, 1332 pages ISBN 978-1-421438-14-6.

28. Souza RP, Petrella S, Coimbra TL, Maeda AY, Rocco IM, Bisordi I, et al. Isolation of yellow fever virus (YFV) from naturally infected *Haemagogus* (*Conopostegus*) *leucocelaenus* (diptera, cukicudae) in São Paulo State, Brazil, 2009. Rev Inst Med Trop Sao Paulo. 2011;53(3):133-9.

29. Lira-Vieira AR, Gurgel-Gonçalves R, Moreira IM, Yoshizawa MA, Coutinho ML, Prado PS, et al. Ecological aspects of mosquitoes (Diptera: Culicidae) in the gallery forest of Brasília National Park, Brazil, with an emphasis on potential vectors of yellow fever. Rev Soc Bras Med Trop. 2013;46(5):566–74.

30. Guimarães AE, Lopes CM, de Mello RP, Alencar J. [Mosquito (Diptera, Culicidae) ecology in the Iguaçu National Park, Brazil: 1 Habitat distribution]. Cad Saude Publica. 2003;19(4):1107-16.

31. Müller GA, Kuwabara EF, Duque JE, Navarro-Silva MA, Marcondes CB. New records of mosquito species (Diptera: Culicidae) for Santa Catarina and Paraná (Brazil). Biota Neotrop. 2008; 8(4).

32. Goenaga S, Fabbri C, Dueñas JC, Gardenal CN, Rossi GC, Calderon G, et al. Isolation of yellow fever virus from mosquitoes in Misiones province, Argentina. Vector Borne Zoonotic Dis. 2012 Nov;12(11):986–93.

33. Couto-Lima D, Madec Y, Bersot MI, Campos SS, Motta MA, Santos FBD, et al. Potential risk of re-emergence of urban transmission of Yellow Fever virus in Brazil facilitated by competent *Aedes* populations. Sci Rep. 2017 Jul 7;7(1):4848.

34. Rezende IM, Sacchetto L, Munhoz de Mello É, Alves PA, Iani FCM, Adelino TÉR, et al. Persistence of Yellow fever virus outside the Amazon Basin, causing epidemics in Southeast Brazil, from 2016 to 2018. PLoS Negl Trop Dis. 2018 Jun;12(6):e0006538.

